# A Bile Duct-on-a-Chip with Organ-Level Functions

**DOI:** 10.1101/594291

**Authors:** Yu Du, Gauri Khandekar, Jessica Llewellyn, William Polacheck, Christopher S. Chen, Rebecca G. Wells

**Affiliations:** Division of Gastroenterology, Department of Medicine, Perelman School of Medicine at the University of Pennsylvania, Philadelphia, PA; The Wyss Institute for Biologically Inspired Engineering, Harvard University, Boston, MA; The Biological Design Center and Department of Biomedical Engineering, Boston University, Boston, MA; Joint Department of Biomedical Engineering, University of North Carolina at Chapel Hill and North Carolina State University, Chapel Hill, NC; Tissue Microfabrication Laboratory, Department of Biomedical Engineering, Boston University, Boston, MA 02215; Department of Bioengineering, School of Engineering and Applied Sciences, The University of Pennsylvania, Philadelphia, PA; Department of Pathology and Laboratory Medicine, Perelman School of Medicine at the University of Pennsylvania, Philadelphia, PA; Center for Engineering MechanoBiology, The University of Pennsylvania, Philadelphia, PA

**Keywords:** Microfluidics, glycocalyx, biliatresone, permeability, monolayer

## Abstract

Chronic cholestatic liver diseases such as primary biliary cholangitis (PBC) and primary sclerosing cholangitis (PSC) are frequently associated with damage to the barrier function of the biliary epithelium, but barrier function is difficult to study in vivo and has not been recapitulated in vitro. Here we report the development of a bile duct-on-a-chip that phenocopies not only the tubular architecture of the bile duct in three dimensions, but also its barrier functions. We demonstrated that mouse cholangiocytes in the channel of the device became polarized and formed mature tight junctions, and that the permeability of the cholangiocyte monolayer was comparable to that measured ex vivo for the rat bile duct. Permeability decreased significantly when cells formed a compacted monolayer with cell densities comparable to that seen in vivo. This device enabled independent access to the apical and basolateral surfaces of the cholangiocyte channel, allowing proof-of-concept toxicity studies with the biliary toxin biliatresone and the bile acid glycochenodeoxycholic acid. The cholangiocyte basolateral side was more vulnerable than the apical side to treatment with either agent, suggesting a protective adaptation of the apical surface that is normally exposed to bile. Further studies revealed a protective role of the cholangiocyte apical glycocalyx, wherein disruption of the glycocalyx with neuraminidase increased the permeability of the cholangiocyte monolayer after treatment with glycochenodeoxycholic acid. *Conclusion*: This bile duct-on-a-chip captured essential features of a simplified bile duct in structure and organ-level functions and represents a novel in vitro platform to study the pathophysiology of the bile duct using cholangiocytes from a variety of sources.

## Introduction

Chronic cholestatic liver diseases such as primary biliary cholangitis (PBC) and primary sclerosing cholangitis (PSC) are often associated with alterations in the tight junctions of bile duct epithelial cells (cholangiocytes). Data from mouse models similarly suggest that impaired tight junction integrity plays an important role in the pathogenesis of cholangiopathies^(1-3)^. Most in vitro research on biliary physiology and pathology, however, employs cells either cultured in 2D monolayers or as organoids in 3D extracellular matrix (ECM). These conventional methods fail to replicate many key aspects of bile duct structural organization or to recapitulate important tissue-level integrated physiological functions such as forming a protective barrier and compartmentalizing bile.

Microfluidic organs-on-chips can overcome some of these limitations^(4)^. Soft lithography, which is used to design organs-on-chips, allows control of surface features over a range of physiologically-relevant scales. Organs-on-chips can mimic aspects of the physiology of cell-cell and cell-ECM junctions in tissues including the alveolar-capillary interface^(5)^, the blood-brain barrier^(6-8)^, and liver sinusoids, in a controllable way, allowing both high-resolution imaging and biochemical and metabolic analyses in real time. The technology has great potential to advance the study of tissue development, physiology and pathophysiology.

To capture the structure and environment of the bile ducts, we used organ-on-chip technology to develop a microengineered bile duct with controllable architecture and surrounding matrix. We demonstrate that a self-organized cholangiocyte-lined channel faithfully recapitulates key functions of the bile duct and enables us to study the barrier function of cholangiocytes monolayer quantitatively and independently from either the apical or basolateral side. As a proof-of-concept, we use this model to demonstrate the protective role of the cholangiocyte apical glycocalyx.

## Materials and Methods

### CELL ISOLATION AND CULTURE

The small cholangiocyte cell line was originally isolated from normal mice (BALB/c) and immortalized by transfection with the SV40 large-T antigen^(9)^. Primary cholangiocytes were isolated from extrahepatic bile ducts (EHBD) of normal adult mice (BALB/c) as described previously^(10)^. Cells were cultured in low glucose Dulbecco’s Modified Eagle Medium (DMEM; Thermo Fisher Scientific, Waltham, MA) supplemented as previously reported^(9-11)^.

### FABRICATION OF THE BILE DUCT-ON-A-CHIP

Microfluidic devices were fabricated using soft lithography as described previously^(12, 13)^ (Fig. 1A). Polydimethylsiloxane (PDMS, Sylgard 184, Dow-Corning, Midland, MI) devices were treated with 0.01% (v/v) poly-L-lysine for 1 h and 0.5% (v/v) glutaraldehyde for 20 min to promote collagen adhesion. After the devices were washed overnight in water and for 30 min in 70% ethanol, steel acupuncture needles (160 μm diameter, Seirin, Kyoto, Japan) were inserted and the devices were then sterilized under UV for 20 min. A solution of 2.5 mg/ml rat tail type 1 collagen (Thermo Fisher Scientific), 1X DMEM medium, 10 mM HEPES, 0.1 M NaOH and NaHCO_3_ (0.035% w/v) was infused via the side ports and allowed to polymerize for 20 min at 37°C. Needles were removed to form channels, which were then coated with 100 μg/ml laminin overnight (via the large reservoir ports) at 37°C. A suspension of 0.5 million/ml cholangiocytes was introduced into the reservoir ports. Cells were allowed to adhere to the top surface of the channel for 2 min; devices were then flipped to allow cells to adhere to the bottom surface of the channel for 5 min. Non-adherent cells were removed by rinsing with cell culture medium and the devices were filled with fresh medium. Devices were maintained at 37°C (5% CO_2_) for 7 d on a rocker at 5 rpm with daily medium changes until the development of compact monolayers.

**Figure 1.**
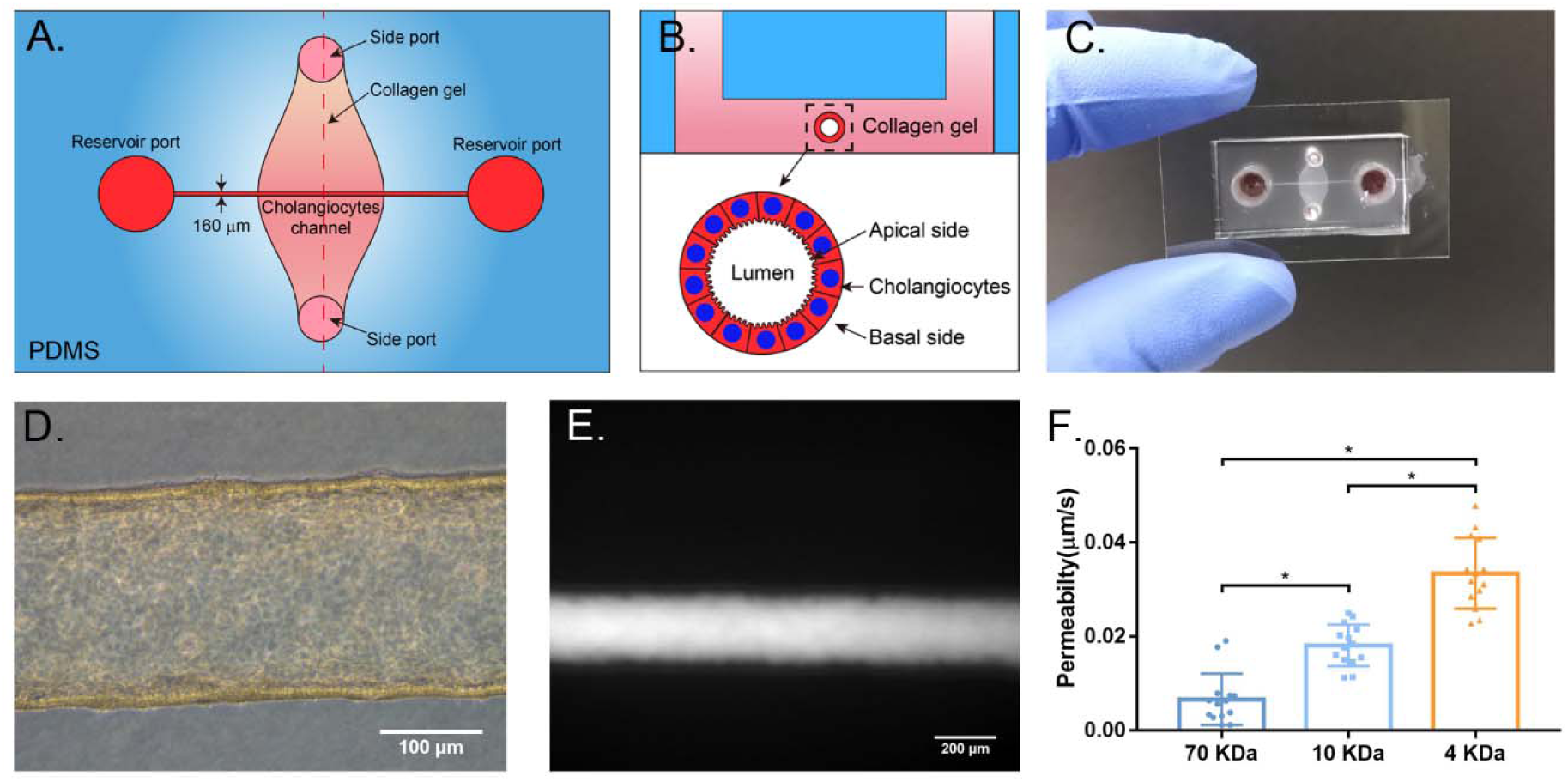
Fabrication and characterization of a three dimensional biomimetic bile duct-on-a-chip. A. Schematic of top view of the bile duct-on-a-chip. B. Schematic of the device in cross section. C. Image of an actual bile duct-on-a-chip, top view. D. Representative bright field image of the channel lined by a layer of mouse cholangiocytes (cell line). Scale bar, 100 μm. E. Representative image of FITC-dextran (70 kDa) in the channel, imaged after 10 min. Scale bar, 200 μm. F. Permeability of the cholangiocyte (cell line)-lined channel to FITC-dextran (70 kDa, 10 kDa and 4 kDa), n=14 devices, each device tested sequentially with FITC-dextran from 70 to 4 kDa. All data are presented as mean ± SD, *P<0.05.

### PERMEABILITY MEASUREMENTS

To measure the permeability of the cholangiocyte monolayers in the device, fluorescent dextran (70 kDa, 10 kDa and 4 kDa, labeled with fluorescein isothiocyanate (FITC) (Sigma, St. Louis, MO)) in phosphate buffered saline (PBS) was introduced into the devices at a concentration of 20 μg/ml. Diffusion of the dextran was imaged in real time with an EVOS FL Auto 2 Imaging System (Thermo Fisher Scientific) at 10X magnification. The diffusive permeability coefficient (Pd) was calculated by measuring the flux of dextran into the collagen gel and fitting the resulting diffusion profiles to a dynamic mass conservation equation, as described previously^(12, 14)^.

### IMMUNOSTAINING AND LECTIN STAINING

Cholangiocyte monolayers in the device were fixed with 4% Paraformaldehyde (PFA) at 37°C for 20 min with rocking. Cells were rinsed 3X with PBS and permeabilized with 0.1% Triton X-100 for 3 d, then blocked with 2% bovine serum albumin (BSA) (Sigma) in PBS at 4°C overnight with rocking. Primary antibodies together with 4′,6-diamidino-2-phenylindole (DAPI; Thermo Fisher Scientific) diluted in 2% BSA in PBS were incubated overnight at 4°C, and rinsed 3X with PBS for 5 min each with rocking, followed by an overnight rinse. Secondary antibodies were diluted in 2% BSA in PBS and incubated overnight at 4°C, and rinsed 3X with PBS for 5 min each on a rocker, followed by an overnight rinse. Primary antibodies and the concentrations used are listed in Supplemental Table S1. Cy3- and Cy5-conjugated secondary antibodies were used at 1:400 (Vector Laboratories, Burlington, CA).

Extrahepatic bile ducts were frozen in O.C.T. (Tissue Tek, VWR, Bridgeport, NJ). 5 µm thick sections were brought to room temperature and fixed in 10% neutral buffered formalin for 4 min. Tissue was blocked with StartingBlock™ T20/PBS Blocking Buffer (Thermo Scientific) before being incubated overnight with K19 primary antibody (1:100, Troma III, DSHB, Iowa city, Iowa) in staining buffer (0.2% Triton X-100, 0.1% bovine serum albumin, in PBS) at 4°C. Cy™2 AffiniPure Donkey Anti-Rat IgG (H+L) (1:500, Jackson ImmunoResearch Lab, West Grove, PA) was then used as the secondary antibody and incubated for 1 hr at RT.

For Sambucus nigra (SNA) and soybean agglutinin (SBA) lectin staining, cholangiocytes were fixed with 4% PFA at 37°C for 20 min with rocking. Cells were rinsed with PBS for 1 h, then blocked with 1X Carbo-Free Blocking Solution (Vector Laboratories) for 1 h at RT on a rocker. After 3X 15 min PBS rinses, cells were incubated with SBA-FITC or SNA-Cy5 (Vector Laboratories) at 20 μg/ml for 1 h at RT with rocking. Cells were rinsed with PBS for 1 h, then the channel was filled with mounting medium containing DAPI (Vector Laboratories). Images were acquired using a SCTR Leica confocal microscope and Leica application suite (LAS X) (Leica, Buffalo Grove, IL).

### STATISTICAL ANALYSIS

Statistical significance was assessed using one-way ANOVA. p<0.05 was regarded as statistically significant and calculated with Prism 7 (GraphPad Software, La Jolla, CA). All data are presented as mean ± standard deviation (SD). Sample size is indicated in the corresponding figure legends.

## Results

### FABRICATION OF THE BILE DUCT-ON-A-CHIP

In order to study the function of the apical and basal sides of cholangiocytes independently, we developed a microfluidic device consisting of a PDMS support with a 160 μm cholangiocyte-lined channel through a matrix bulk; apical and basal sides of the channel were accessible through separate ports (Figure 1A, B). The device was designed as per previously-described models of endothelium-lined, vascular-mimetic channels^(12, 13)^. To generate a bile duct-on-a-chip, the channel was formed within a collagen plug and the lumen was coated with laminin (100 μg/ml) for 4 hours. The channels were seeded with a line of mouse cholangiocytes. Cholangiocytes in the channel formed a confluent and then compact epithelial monolayer (nearly cubical cells with similar height and width) (Figure 1C, D).

To determine whether the bile duct-on-a-chip recapitulated the barrier function of the bile duct, we perfused the lumen with FITC-dextran ranging in size from 4 to 70 kDa. There was no obvious leakage of fluorescent dextran into the collagen matrix even after 10 minutes (Figure 1E). Quantification of permeability showed an increase as the molecular weight of the dextran decreased (Figure 1F). The measured permeability of cholangiocyte cell line was comparable with that measured in rat ex vivo systems using insulin (5.8 kDa; 0.45 μm/s)^(15)^. This model of the bile duct thus mimics the barrier function of ducts in vivo.

### CHARACTERIZATION OF THE BILE DUCT-ON-A-CHIP

We used immunofluorescence staining to characterize the cholangiocytes in the channel. F-actin staining showed that cholangiocytes grew into a compact monolayer, forming a cylindrical tube with two open ends connected to the large reservoirs (Figure 2A, C). Cells maintained expression of the cholangiocyte marker K19 (Figure 2B), and demonstrated staining for ZO-1 (TJP-1) and E-cadherin 1 (CDH1, also known as cadherin 1) at cell-cell junctions (Figure 2D, E), confirming the formation of tight junctions. Staining for the apical sodium-dependent bile acid transporter (ASBT) confirmed that cells in the bile duct-on-a-chip were polarized (Figure 2F).

**Figure 2.**
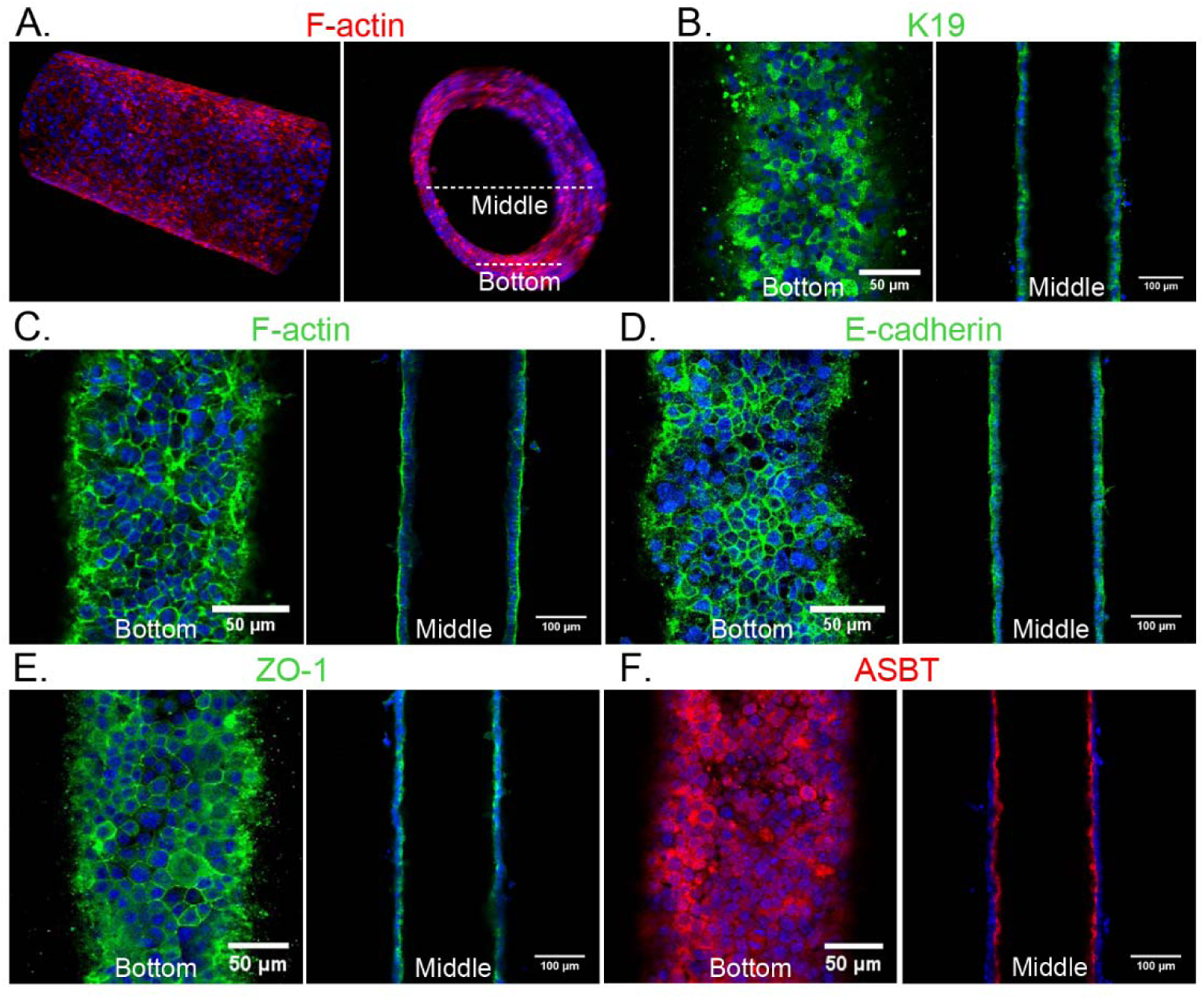
Characterization of a channel lined with mouse cholangiocytes (cell line). A. Representative confocal images of cholangiocytes in the bile duct-on-a-chip forming a monolayer within the cylindrical channel, stained for F-actin (red) and nuclei (DAPI; blue). Longitudinal (left panel) and cross-sectional (right panel) views. White dashed lines indicate the bottom surface and cross section through the center shown in the remaining panels. B-F. Immunofluorescent images across the bottom (left panels in B-F) and middle (right panels in B-F) of the channel stained with antibodies against (B) K19, (C) F-actin, (D) E-cadherin, E) ZO-1, and (F) the apical sodium bile salt transporter (ASBT). Nuclei shown by DAPI staining (blue). Images are representative of at least independently constructed devices for each condition. Scale bars: 50 μm, left panels; 100 μm right panels.

### A SUPERCONFLUENT MONOLAYER IS REQUIRED FOR OPTIMAL BARRIER FUNCTION

Full barrier function required that cholangiocytes were not just confluent but “superconfluent” and compact (Figure 3A); as the cholangiocyte monolayer grew more dense, permeability decreased significantly (Figure 3B). By F-actin staining, we confirmed that the permeability difference between what we term a confluent versus a compact monolayer was not the result of gaps in the monolayer (Figure 3C). Cell height increased more than two-fold in the compact compared to the confluent monolayer (Figure 3D). To determine the cell density of cholangiocyte monolayers in vivo, we counted cell number as a function of monolayer length in adult mouse EHBD and in confluent and compact devices (Figure 3E). We found that compact monolayers were more dense than confluent ones, and that mice EHBD (Figure 3E, F) had an even higher cell density. These results suggested that the biliary epithelium forms a dense, compact monolayer – beyond confluence – in order to form a mature barrier.

**Figure 3.**
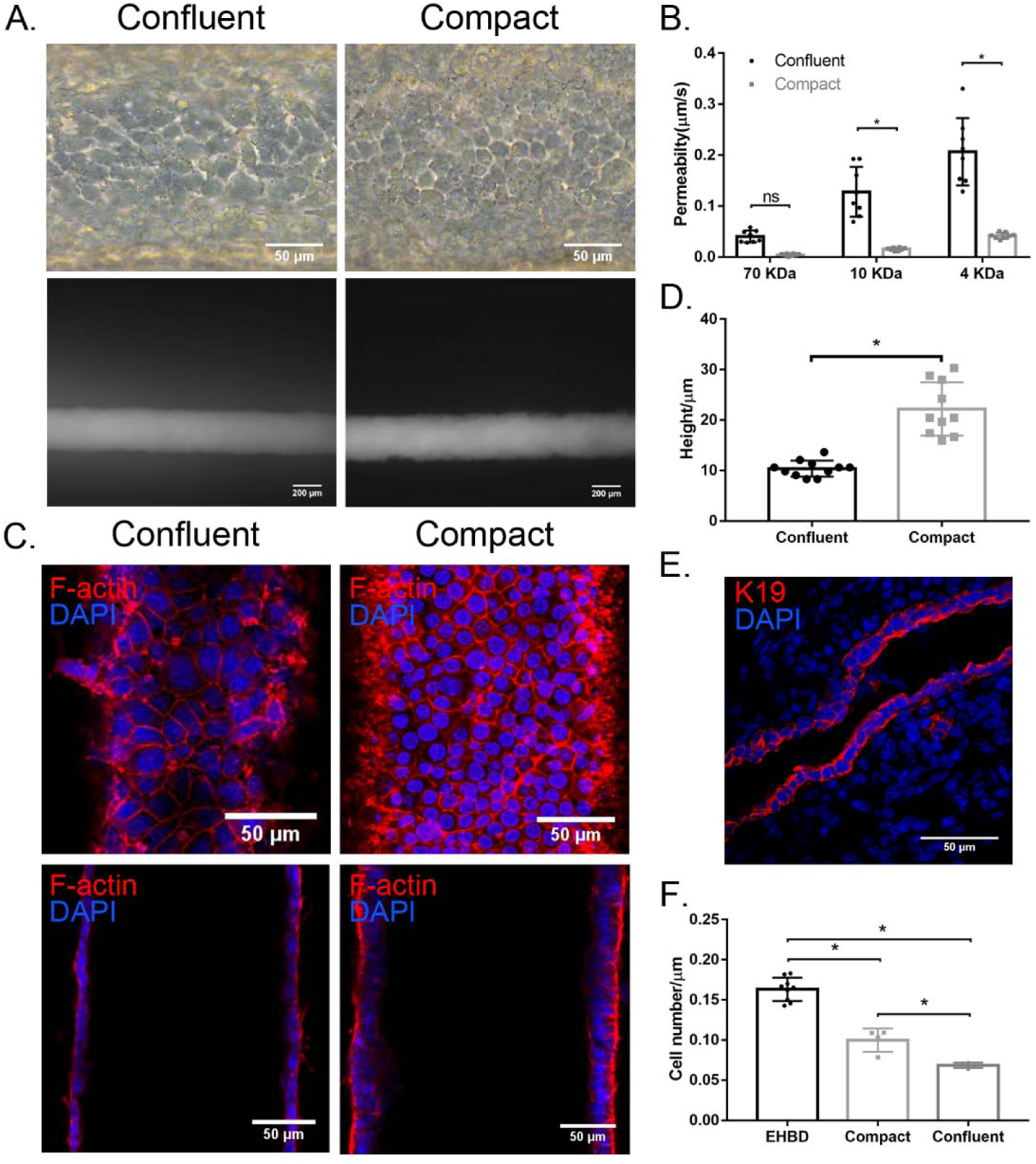
Cholangiocyte monolayers require high confluency for mature barrier function. A. Representative bright field images (upper panels) and images after FITC-dextran (4 kDa) perfusion for 2 min (lower panels) of confluent and compact cholangiocyte channels. Scale bar: 100 μm (upper panels), 200 μm (lower panels). B. Permeability of confluent and compact cholangiocyte channel to FITC-dextran (70 kDa, 10 kDa and 4 kDa), n= 8 devices. C. Bottom (upper panels) and middle (lower panels) views of confluent and compact cholangiocyte monolayers in the devices, stained for F-actin (red) and nuclei (DAPI; blue). Scale bars, 50 μm. D. Cell height of confluent and compact cholangiocyte monolayers in the devices, n≥ 10. E. Representative images of adult mouse extrahepatic bile duct, stained for K19 (red) and nuclei (DAPI; blue), n=9. Scale bar, 50 μm. F. Cell density in confluent, compact cholangiocyte channel and mice extrahepatic bile duct, n= 4-9. Images are representative of at least three independent experiments. All data are presented as mean ± SD, *P<0.05.

### THE APICAL AND BASAL SURFACES OF THE MONOLAYER CAN BE ACCESSED INDEPENDENTLY

In order to determine whether the apical and basal surfaces of cholangiocytes in the device could be accessed independently, we applied FITC-dextran through either the large reservoir ports, which are directly connected to the lumen, or the small side ports, which connect to the basal surface of the monolayer through the collagen bulk. We found that FITC-dextran added through the reservoir ports remained within the lumen (Figure 4A, B), making contact only with the apical side of the cholangiocytes in the device and unable to penetrate through the monolayer. Dextran applied via the side ports remained within the collagen plug, in contact with only the basal side of the monolayer (Figure 4C, D).

**Figure 4.**
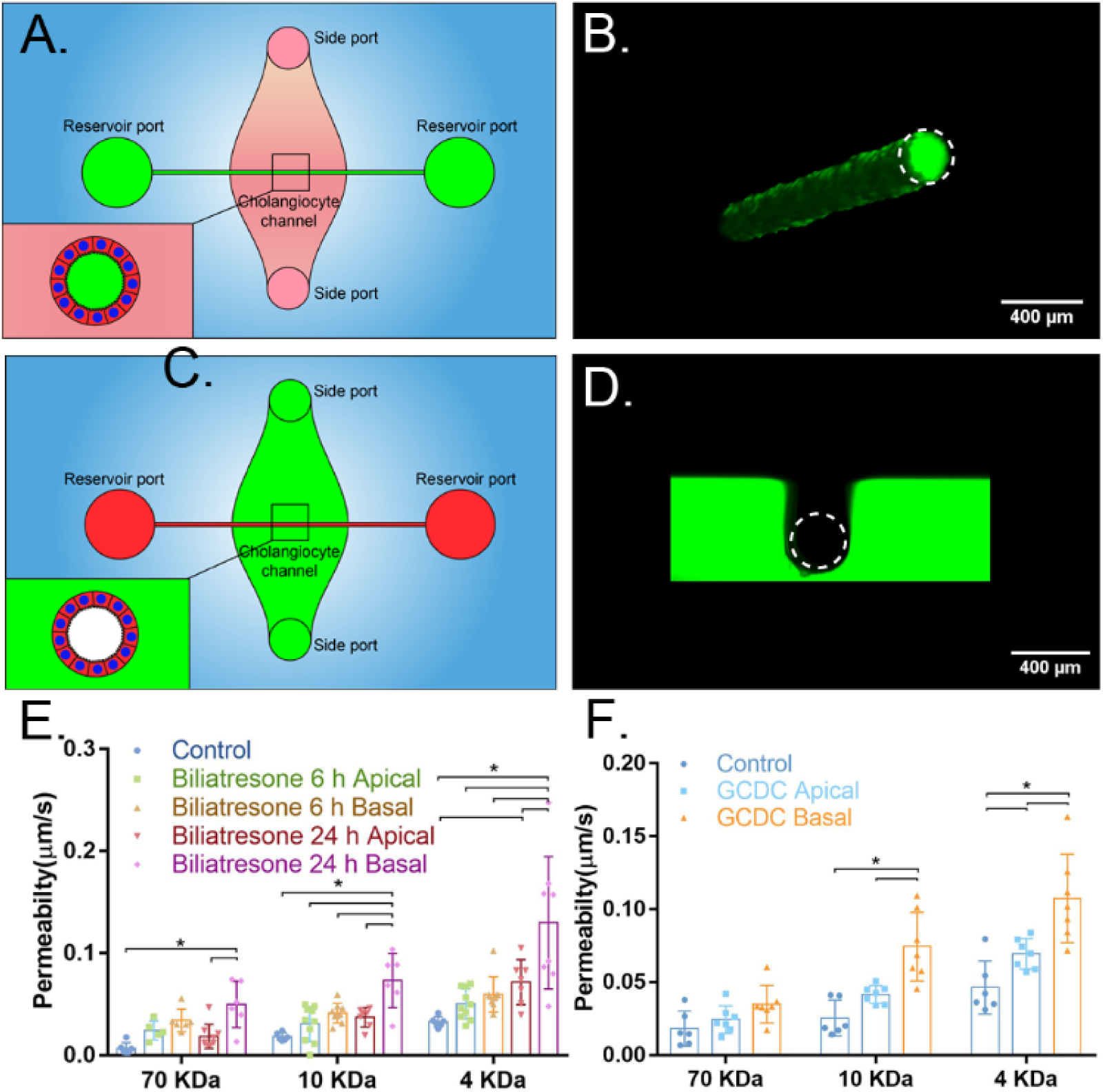
The apical and basal surfaces of cholangiocytes can be treated independently. A. Schematic showing access to the apical side of cholangiocytes through the reservoir ports. B. Confocal image of FITC-dextran solution (green) in the lumen after administration through the reservoir ports. Scale bar, 400 μm. C. Schematic showing access to the basal side of cholangiocytes through the side ports. D. Confocal image of FITC-dextran solution (green) in the collagen bulk and surrounding but not within the lumen after administration through the side ports. Scale bar, 400 μm. E. Diffusive permeability across monolayers treated with biliatresone via apical or basal surfaces for 6 h or 24 h, as measured using FITC-dextran (70 kDa, 10 kDa and 4 kDa) in the lumen, n ≥ 6 devices for each condition. F. Diffusive permeability across monolayers treated with 1 mM GCDC via apical or basal surfaces for 1 h, as measured using FITC-dextran (70 kDa, 10 kDa and 4 kDa) in the lumen, n ≥ 6 devices for each condition. All data are presented as a mean ± SD, *P<0.05.

Biliatresone is a plant isoflavonoid that is toxic to cholangiocytes and causes a syndrome mimicking the pediatric disease biliary atresia in neonatal livestock and larval zebrafish^(16)^. We previously demonstrated that biliatresone applied to the basal surface of a cholangiocyte spheroid causes mislocalization of the apical markers ZO-1 and E-cadherin and increases cholangiocyte monolayer permeability^(11)^. A limitation of these spheroid studies was that biliatresone could only be added to the basal surface. As a demonstration of the potential use of the bile duct-on-a-chip device, we showed that biliatresone treatment caused cholangiocyte damage, as evidenced by increased permeability of monolayers in the device, and that this damage was worse with basal as opposed to apical administration (Figure 4E).

Cholangiocytes are adapted to tolerate exposure to toxic bile at their apical surfaces though it has been difficult to study in cell culture. Impaired duct barrier function may lead to bile leakage through the monolayer, and cause damage from the basolateral side, which is less tolerant of bile ^(17-19)^. As a second illustration of the potential use of the bile duct-on-a-chip device, we exposed cholangiocytes to 1 mM of the bile acid glycochenodeoxycholic acid (GCDC) from either the apical or basal side. GCDC treatment caused significantly more damage, as defined by increased permeability, when applied to the basal side rather than the apical side (Figure 4F), suggesting that once bile leaks through the epithelial monolayer, it can cause additional damage through the basolateral side of the barrier, leading to a feedback loop of further damage and increased leakage.

### THE GLYCOCALYX PROTECTS CHOLANGIOCYTES FROM BILE ACID-INDUCED DAMAGE

Cholangiocytes have an apical glycocalyx that protects them from bile acid toxicity^(20, 21)^. We stained the apical surfaces of the cholangiocyte monolayer in the bile duct-on-a-chip using the lectin Sambucus nigra agglutinin (SNA), which binds to α_2-6_-bound (terminal) sialic acid residues, and demonstrated that the sialic acid-enriched glycocalyx was maintained in cholangiocytes in the device and was appropriately localized to the apical surface (Figure 5A). We also stained cells in the channel with the lectin soybean agglutinin (SBA), which only detects carbohydrates *lacking* terminal sialic acid residues (Figure 5E), confirming a lack of cholangiocyte staining at baseline (Figure 5B). We then treated cells with neuraminidase, applied apically, to remove the terminal sialic acid. Post-treatment staining with SNA was minimal, while there was new staining with SBA, confirming the presence of apical sialic acid residues in untreated cells and the efficacy of neuraminidase in this system (Figure 5C, D). To test the role of the glycocalyx in protecting cholangiocytes from bile acids, we perfused the device lumen with GCDC (1 mM) before or after treatment with neuraminidase, and then measured monolayer permeability. We found no effect of GCDC, with or without neuraminidase treatment, on permeability to 70 kDa FITC-dextran; however, GCDC alone increased the permeability to 4 kDa FITC-dextran, and GCDC exposure of neuraminidase-treated cholangiocytes increased permeability to both 4 kDa and 10 kDa FITC-dextran (Figure 5F). Taken together, these findings demonstrate that the cholangiocyte-lined channel is resistant to bile acid toxicity and that the glycocalyx plays a protective role.

**Figure 5.**
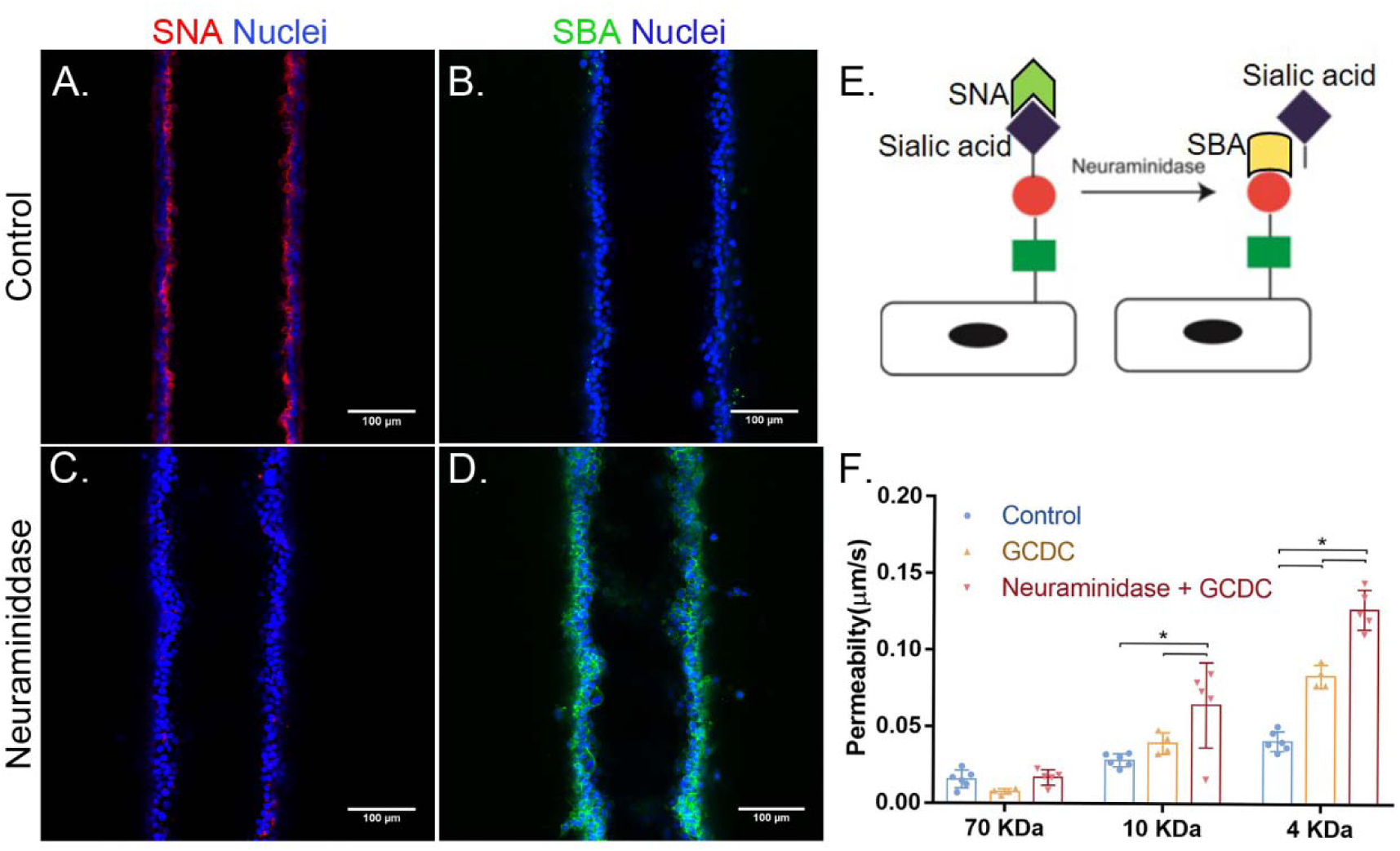
An intact glycocalyx protects cholangiocytes from bile acids. A-D. Staining using the lectins SNA and SBA in the bile duct-on-a-chip, (A, B) before and (C, D) after neuraminidase treatment. Scale bars, 100 μm. E. Schematic showing that SNA recognizes sialyated carbohydrates, and that removal of sialic acid is required for recognition by SBA. F. Diffusive permeability of devices treated with the bile acid GCDC with or without prior desialyation with neuraminidase, n=4-6 devices for each condition. All data are presented as a mean ± SD, *P<0.05.

### CONSTRUCTION OF A BILE DUCT-ON-A-CHIP WITH PRIMARY MURINE EXTRAHEPATIC CHOLANGIOCYTES

To demonstrate that the bile duct-on-a-chip can be used with cells from various sources, we also used primary murine extrahepatic cholangiocytes (Figure S1A). Primary cholangiocytes, like the cell line, formed a compact tubular monolayer. Cells maintained expression of the cholangiocyte marker K19 (Figure 6A), and demonstrated staining for E-cadherin and ZO-1 at cell-cell junctions (Figure 6C, D), confirming the formation of tight junctions. Staining for ASBT (Figure 6B) and acetylated α-tubulin (Figure 6E, F) confirmed that primary cholangiocytes in the bile duct-on-a-chip were ciliated and highly polarized. In permeability assays, there was no visible leakage of 4kDa FITC-dextran from the lumen even after 10 mins (Figure S1B, Movie S1). Analysis of the intensity change along a line perpendicular to the channel showed almost no increase in intensity across the gel after 2 mins (Figure S1D), in contrast to the cholangiocyte cell line (Figure S1C) and the permeability to 4 kDa FITC-dextran of primary cholangiocyte was about only 16% of that for the cholangiocyte cell line (Figure S1E). Thus, the primary cell-lined device demonstrated even better barrier function than the device lined with the cholangiocyte cell line (Figure S1E).

**Figure 6.**
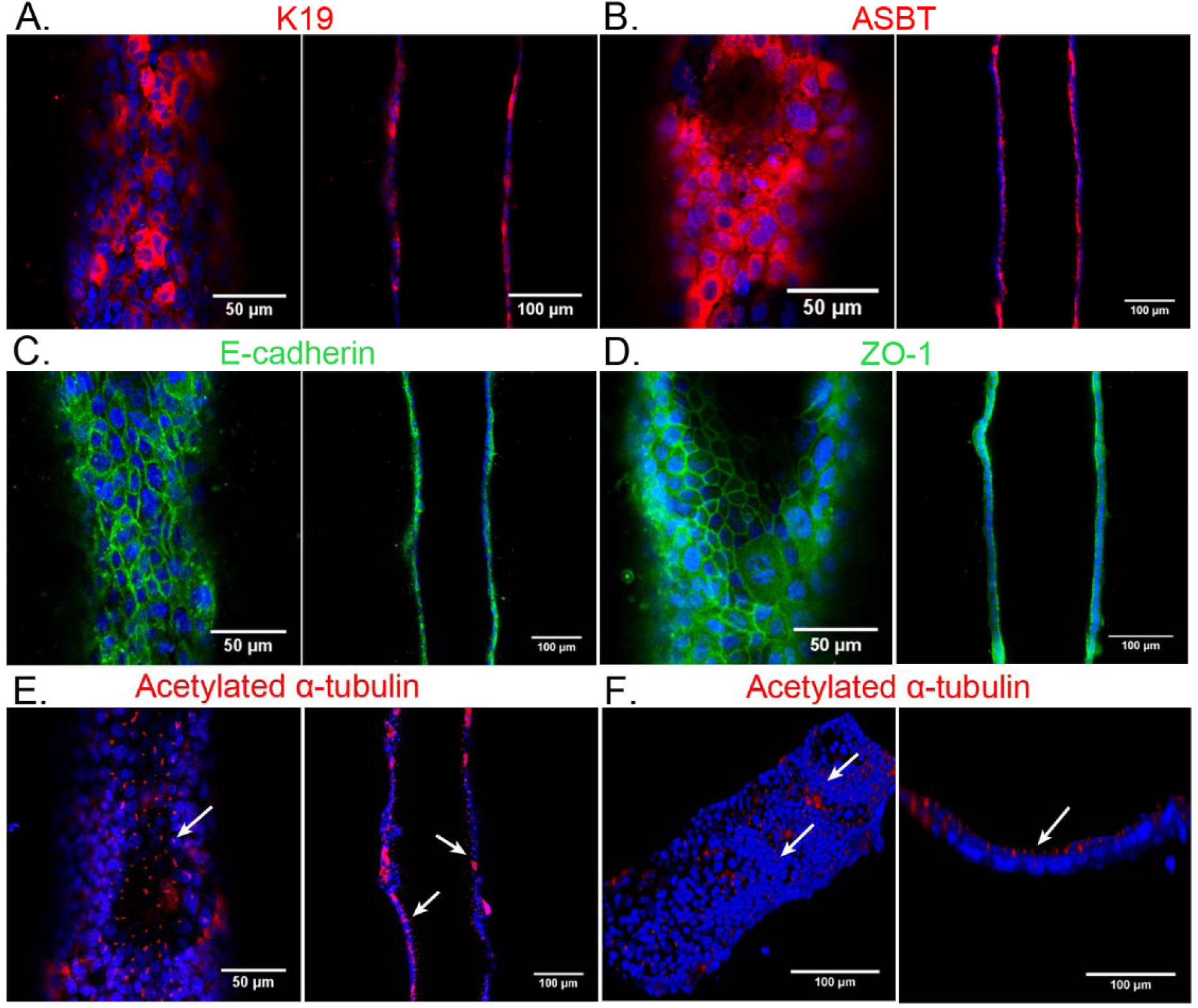
Bile duct-on-a-chip with primary murine extrahepatic cholangiocytes. Immunofluorescence images across the bottom (left panels) and middle (right panels) of channels stained with antibodies (shown in red or green) against (A) K19, (B) ASBT, (C) E-cadherin, (D) ZO-1, and (E,F) acetylated α-tubulin, with DAPI nuclear staining (blue). Top (left panel F) and cross-sectional (right panel F) views of cilia (white arrow) in the cholangiocyte channel. Scale bars, 50 μm (left panels except F); 100 μm (right panels and F, left panel). Images are representative of three independent experiments.

## Discussion

We have developed an in vitro model of the bile duct by employing organ-on-chip technology, recapitulating not only the three-dimensional architecture of the bile duct but also its barrier function. We demonstrated that cholangiocytes are polarized in the device and that permeability is comparable to in vivo values for the bile duct, requires a dense monolayer, and can be quantified. We also showed that the device enables apical vs. basolateral treatments and that cells respond differently depending on the side of exposure. As a demonstration of the potential use of the device in studying biliary physiology, we showed that the apical glycocalyx plays a protective role against bile acid toxicity and that the biliary toxin biliatresone has greater toxicity when applied to the basal surface. The bile duct-on-a-chip thus represents a novel in vitro model for studying cholangiopathies. The in vivo-like permeability barrier and the potential to quantify changes in barrier function particular strengths of this model.

Most in vitro studies of cholangiopathies have relied on 2D cell culture or on 3D organoid culture. 2D culture, although convenient, fails to replicate bile duct physiology with respect to duct structure, extracellular matrix composition, and stiffness, and does not support fluid flow. Polarized cell organization and permeability measurements, although possible, require specialized culture systems. Cholangiocytes in 3D spheroid culture differentiate well, can be cultured in physiological matrices, and demonstrate barrier functions and transport activities. However, spheroids are highly variable in shape and size, and it is difficult to control their positions in a gel (for microscopy), sample lumen content, or treat the cholangiocyte apical surface. The bile duct-on-a-chip provides uniform ducts that are accessible for imaging, with two sets of ports that enable sampling of luminal contents as well as selective (apical vs. basolateral) exposures. Additionally, the chemical composition and mechanics of the surrounding matrix as well as the fluid flow rate can be varied.

Several different bioengineered bile ducts, including acellular constructs using biodegradable materials and constructs using human cholangiocytes, have been reported. Sampaziotis et al. reconstructed the biliary epithelium with a bioengineered bile duct consisting of extrahepatic cholangiocytes derived from the common bile duct or the gallbladder. These results suggested that bioengineered bile ducts may have promise for transplantation applications, but the difficulty in manipulating this duct construct makes it unsuitable for in vitro study. Chen et al. constructed a bile duct using organoid-derived cholangiocyte-like cells on a collagen-coated polyethersulfone hollow fiber membrane, yielding a tubular structure with polarized bile acid transport activity. However, the cholangiocytes cultured on the fiber have outward-facing apical surfaces, in contrast to ducts in vivo and the bile duct-on-a-chip.

Previous studies have suggested that epithelial cell confluence is associated with morphology, polarity and barrier function^(22-25)^. Although changes in transepithelial electronic resistance have been reported to be minimal after cells reach confluence^(24)^, we found that permeability decreased significantly when cells progressed from confluent to compact, with associated changes in height, suggesting changes in tight junction molecules^(25)^. These results demonstrate that compactness is required for an optimal barrier function, and suggest that confluence may be an intermediate rather than ultimate goal in the repair of an injured epithelial layer.

We demonstrated that this device can be constructed with primary murine extrahepatic cholangiocytes and that in the device these cells stably expressed biliary markers, junctional molecules and bile salt transport proteins and also developed cilia on their apical surfaces, facing the lumen. The bile duct-on-a-chip thus offers the opportunity to carry out experiments using genetically-modified cholangiocytes and to compare intra- vs. extra-hepatic cholangiocytes. Human cholangiocytes, including those derived from induced pluripotent stem cells, could also be studied using this device.

In conclusion, our bile duct-on-a-chip mimics the basic features of the bile duct, making it a novel platform for in vitro studies of biliary physiopathology.

## Abbreviations

PBC: primary biliary cholangitis
PSC: primary sclerosing cholangitis
ECM: extracellular matrix
EHBD: extrahepatic bile duct
DMEM: Dulbecco’s Modified Eagle Medium
PDMS: polydimethylsiloxane
FITC: fluorescein isothiocyanate
PBS: phosphate buffered saline
Pd: permeability coefficient
PFA: paraformaldehyde
BSA: bovine serum albumin
DAPI: 4′,6-diamidino-2-phenylindole
SNA: Sambucus nigra lectin
SBA: soybean agglutinin
ASBT: apical sodium-dependent bile salt transporter
GCDC: glycochenodeoxycholic acid

## ACKNOWLEDGEMENTS

We are grateful to the UPenn Cell and Developmental Biology Microscopy Core, the Singh Center for Nanotechnology, and the UPenn NIDDK Center for Molecular Studies in Digestive and Liver Disease (NIH-P30-DK050306). The small cholangiocyte cell line was generously provided by Gianfranco Alpini (Texas A&M Health Science Center College of Medicine and Baylor Scott & White Digestive Disease Research Center).

## Figures and Legends

**Figure S1.**
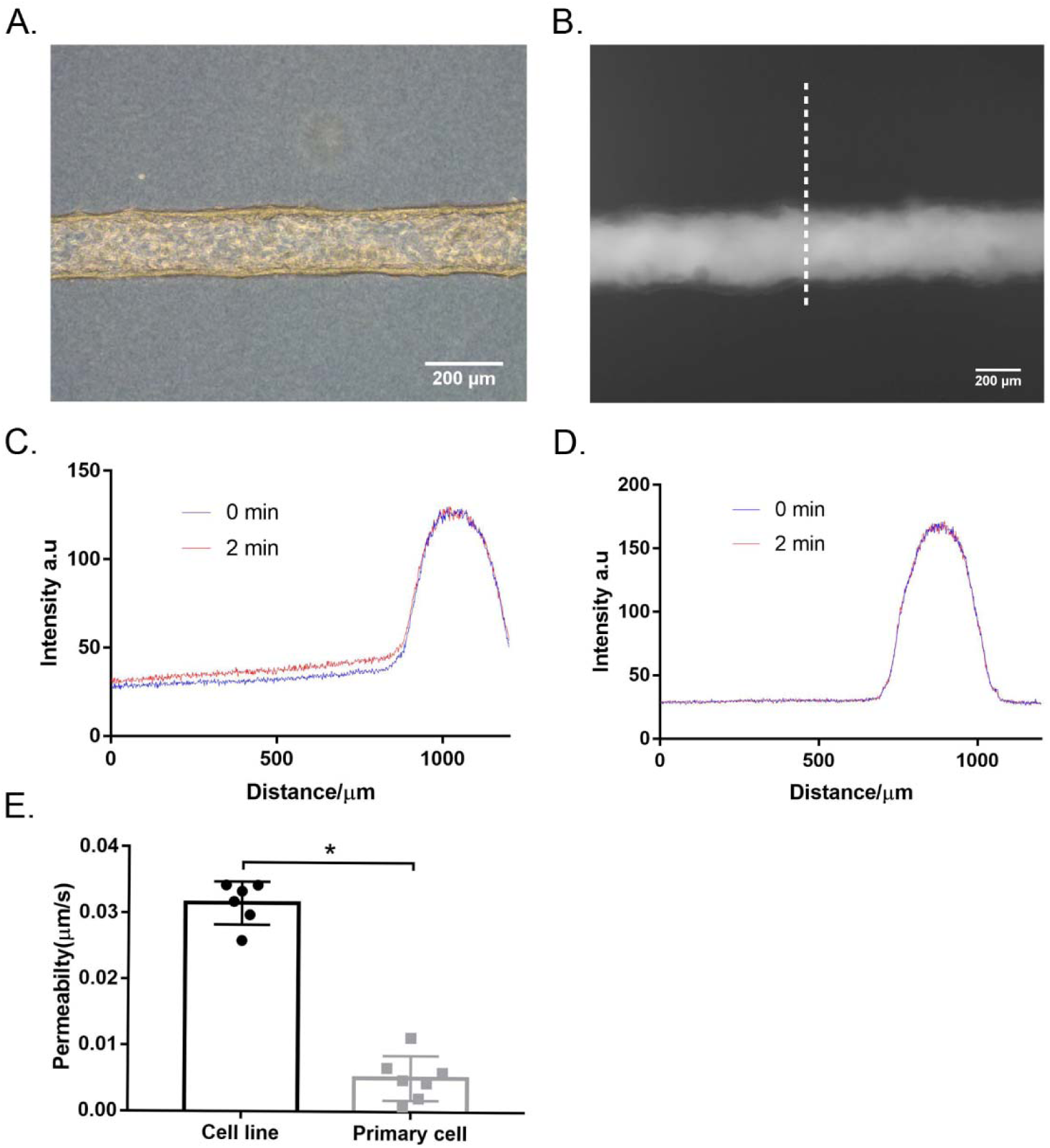
Bile duct-on-a-chip constructed with primary extrahepatic cholangiocytes demonstrated better barrier function than the cell line. A. Representative bright field image of the channel lined by a layer of primary extrahepatic cholangiocytes. B. Representative image of FITC-dextran (4 kDa) in a channel, imaged after 10 min (dashed line used for intensity profile analysis). Scale bars, 200 μm. C. Intensity profile along a line perpendicular to a representative cholangiocyte channel generated using the transformed cholangiocyte cell line at baseline (blue) and 2 min after perfusion with FITC-dextran (4 kDa; red). D. Intensity profile along the dashed line in panel B perpendicular to the cholangiocyte channel (primary extrahepatic cholangiocyte) at baseline (blue) and 2 min after perfusion with FITC-dextran (4 kDa; red). E. Permeability of the cell line and primary cholangiocyte-lined channel to FITC-dextran (4 kDa), n=6-7 devices. All data are presented as mean ± SD, *P<0.05.

Movie S1. Representative time lapse imaging of perfusing FITC-dextran (4 kDa) in the primary cholangiocyte channel for 10 min.

**Supplemental Table S1.**
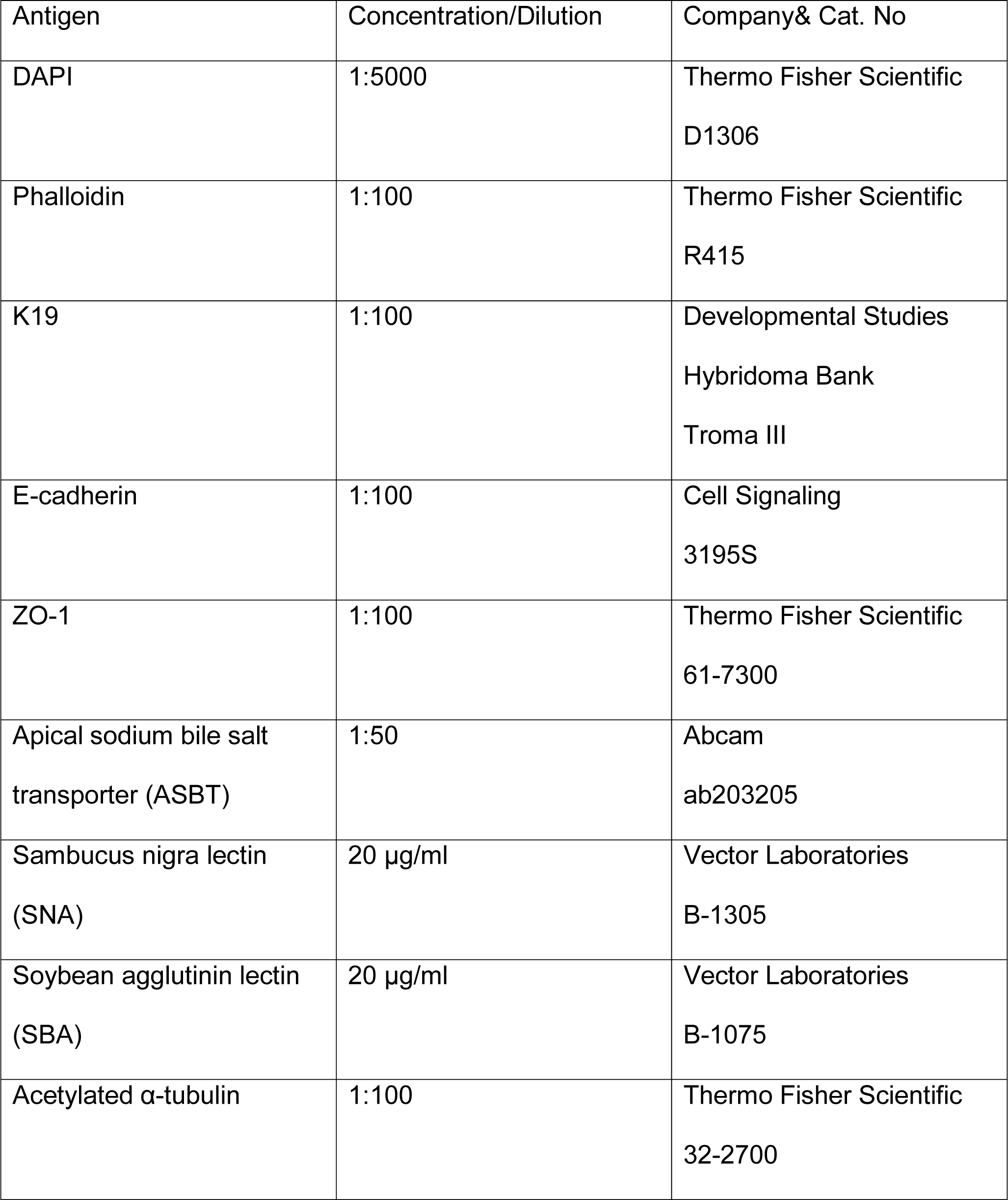

