## Supplemental Table S1 for "A Bile Duct-on-a-Chip with Organ-Level Functions"

| Antigen | Concentration/Dilution | Company& Cat. No |
| --- | --- | --- |
| DAPI | 1:5000 | Thermo Fisher Scientific  D1306 |
| Phalloidin | 1:100 | Thermo Fisher Scientific  R415 |
| K19 | 1:100 | Developmental Studies Hybridoma Bank  Troma III |
| E-cadherin | 1:100 | Cell Signaling  3195S |
| ZO-1 | 1:100 | Thermo Fisher Scientific  61-7300 |
| Apical sodium bile salt transporter (ASBT) | 1:50 | Abcam  ab203205 |
| Sambucus nigra lectin (SNA) | 20 μg/ml | Vector Laboratories  B-1305 |
| Soybean agglutinin lectin (SBA) | 20 μg/ml | Vector Laboratories  B-1075 |
| Acetylated α-tubulin | 1:100 | Thermo Fisher Scientific  32-2700 |
